# Cardiac-specific Kv1.1 deficiency alters cardiomyocyte electrophysiology without modifying overall cardiac function or arrhythmia susceptibility

**DOI:** 10.1101/2025.08.25.671830

**Authors:** Kelsey Paulhus, Man Si, Krystle Trosclair, Ellen Aughenbaugh, Maxine Parkinson, Nicole M. Gautier-Hall, Megan Watts, Frederica Kizek, Md. Shenuarin Bhuiyan, Paari Dominic, Kathryn A. Hamilton, Edward Glasscock

## Abstract

The leading cause of epilepsy-related mortality is sudden unexpected death in epilepsy (SUDEP), resulting from seizure-induced cardiorespiratory arrest by mechanisms that remain unresolved. Mutations in ion channel genes expressed in both brain and heart represent SUDEP risk factors because they can disrupt neural and cardiac rhythms, providing a unified explanation for seizures and lethal arrhythmias. However, the relative contributions of brain-driven mechanisms, heart-intrinsic processes, and seizures to cardiac dysfunction in epilepsy remain unclear. Here, we investigated the heart-specific role of the *Kcna1* gene, which encodes Kv1.1 voltage-gated potassium channel α-subunits expressed in both neurons and cardiomyocytes, where they shape action potential firing and influence seizure and arrhythmia susceptibility. We generated cardiac-specific *Kcna1* conditional knockout (cKO) mice lacking Kv1.1 selectively in cardiomyocytes and assessed their cardiac function using *in vitro* and *in vivo* electrophysiology. Cardiac Kv1.1 deficiency prolonged action potentials in atrial, but not ventricular, cardiomyocytes, demonstrating a direct role for Kv1.1 in atrial repolarization. Despite these cellular effects, cKOs exhibited normal lifespans, electrocardiographic features, heart rate variability, pacing-induced arrhythmia susceptibility, contractility, seizure susceptibility, and seizure-induced mortality. Thus, while loss of cardiac Kv1.1 was sufficient to impair atrial repolarization, it did not reproduce the broader cardiac abnormalities seen in global *Kcna1* knockouts. Given the higher mortality rates of global compared with neural-specific knockouts in our previous studies, cardiac Kv1.1 deficiency, while not lethal alone, may increase vulnerability to seizure-related death when combined with neural deficiency, consistent with a brain-heart dyssynergy that lowers the threshold for fatal events.

## INTRODUCTION

Sudden unexpected death in epilepsy (SUDEP) is the leading cause of epilepsy-related mortality, affecting otherwise reasonably healthy individuals with epilepsy (Nashef *et al*., 2012; Thurman *et al*., 2014). SUDEP typically occurs due to cardiorespiratory arrest in the minutes following a seizure (Ryvlin *et al*., 2013). Mutations in ion channel genes coexpressed in the brain and the heart have been identified as genetic risk factors for SUDEP because they can potentially disrupt both brain and heart rhythms, leading to seizures and lethal cardiac arrhythmias (Glasscock, 2014; Goldman *et al*., 2016; Chahal *et al*., 2020; Yu *et al*., 2023). Although cardiac arrhythmias are commonly associated with seizures, the precise mechanistic relationship between cardiac dysfunction and epilepsy has remained unclear (Ha *et al*., 2025). This is partly because cardiac dysfunction can result from both brain-driven or heart-intrinsic processes (Goh *et al*., 2025). Additionally, the stress of seizures on the heart can lead to structural or electrical remodeling that affects cardiac performance over time (Verrier *et al*., 2020; Surges, 2025; Akyuz *et al*., 2025).

*Kcna1* is an example of a brain-heart ion channel gene linked to SUDEP (Glasscock, 2014). *Kcna1* codes for Kv1.1 voltage-gated potassium channel α-subunits, which are highly expressed in brain axons and to a lesser extent in the heart, where they localize to cardiomyocytes of the atria, ventricles, and sinoatrial node (Glasscock, 2019). Mice lacking Kv1.1 due to global *Kcna1* gene deletion (*Kcna1^-/-^*) are widely used to model SUDEP because they exhibit several key aspects from human SUDEP cases (Glasscock *et al*., 2010; Dhaibar *et al*., 2019). These features include frequent early onset generalized tonic-clonic seizures (GTCS) and seizure-related sudden death associated with cardiorespiratory failure (Smart *et al*., 1998; Dhaibar *et al*., 2019). In addition, more than 20 different *KCNA1* mutations have been found to cause epilepsy in humans and at least one has been linked to a SUDEP case (Klassen *et al*., 2014; Paulhus & Glasscock, 2023).

As a brain-heart channel, Kv1.1 can influence cardiac function via both brain-driven and heart-intrinsic mechanisms. In *Kcna1^-/-^* mice, the global absence of Kv1.1 causes various *in vivo* cardiac abnormalities. Some deficits appear to be neurally mediated, such as increases in the incidence of atrioventricular conduction blocks and heart rate variability, indicative of heightened parasympathetic effects on the heart (Glasscock *et al*., 2010; Mishra *et al*., 2017). During seizures, *Kcna1^-/-^* mice commonly exhibit bradycardia and skipped heartbeats, further suggesting a neural basis (Dhaibar *et al*., 2019; Paulhus *et al*., 2025). Other cardiac abnormalities in *Kcna1^-/-^* mice appear to be more indicative of heart-intrinsic dysfunction. For example, *Kcna1^-/-^* mice exhibit altered arrhythmia susceptibility in intracardiac pacing studies and reduced contractility on echocardiograms (Glasscock *et al*., 2015; Trosclair *et al*., 2021). At the cardiomyocyte level, pharmacological or genetic ablation of Kv1.1 causes cardiac repolarization deficits that lead to prolonged action potentials in the atria, ventricles, and sinoatrial (SA) node (Si *et al*., 2018, 2024; Trosclair *et al*., 2021). In the SA node, the lack of Kv1.1 impairs intrinsic cardiac pacemaking, slowing the firing rates of SA cells and leading to bradycardia when hearts are recorded *ex vivo* apart from the autonomic nervous system (Si *et al*., 2024).

A limitation of these previous studies is that *Kcna1^-/-^* mice have Kv1.1 deleted globally in all tissues which hinders assigning the tissue-specific origins of cardiac dysfunction. Furthermore, the presence of repeated seizures in *Kcna1^-/-^*mice may also alter cardiac function by inducing functional and structural remodeling of the heart (Verrier *et al*., 2020). Therefore, to identify the tissue-specific contributions of the brain to cardiac dysfunction in *Kcna1^-/-^*mice, in a prior study we generated neuron-specific conditional knockout (cKO) mice (Trosclair *et al*., 2020). We found that neuron-specific Kv1.1 deficiency was sufficient to cause SUDEP along with increases in daytime heart rate variability and seizure-associated bradycardia and skipped heartbeats (Trosclair *et al*., 2020). However, the potential cardiac effects of deleting Kv1.1 in the heart, while leaving it intact in the brain, remains unknown.

In this study, we aimed to further dissect the relative contributions of brain-driven mechanisms, heart-intrinsic processes, and seizures to cardiac dysfunction in *Kcna1^-/-^* mice. Specifically, we focused on identifying heart-intrinsic effects due to selective Kv1.1 deficiency in the heart. To do this, we generated *Kcna1* conditional knockout (cKO) mice lacking Kv1.1 only in atrial and ventricular cardiomyocytes, which we refer to herein as cardiac-specific cKO mice. We then tested whether cardiac-specific Kv1.1 deficiency was sufficient to reproduce the cardiac deficits observed in the global *Kcna1* gene deletion model. Using various *in vivo* and *in vitro* electrophysiological techniques, we examined cardiac-specific cKO mice for defects in action potential repolarization, cardiac waveforms, arrhythmia susceptibility, and contractility. We also performed seizure threshold tests to identify potential differences in susceptibility to seizures and seizure-induced mortality. Our data reveal that the primary consequence of cardiac-specific *Kcna1* deficiency is cellular repolarization deficits, primarily in the atria, and that these abnormalities are present in the absence of seizures. These findings further support a fundamental heart-intrinsic role of Kv1.1 in regulating the shape of cardiac action potentials and suggests that Kv1.1 plays a larger functional role in the atria than the ventricles. Additionally, our work provides insight into potential brain-heart dyssynergy that may contribute to SUDEP risk.

## MATERIALS AND METHODS

### Animals

Cardiac-specific *Kcna1* conditional knockout (cKO) mice (i.e., *Myh6*-Cre^+/–^; *Kcna1*^del/–^) were generated by crossing heterozygous *Kcna1* floxed (fl) mice (*Kcna1*^fl/+^) with heterozygous *Kcna1* global knockout (KO) mice (*Kcna1*^+/–^) carrying one copy of the alpha myosin-heavy chain (*Myh6*)-Cre transgene (i.e., *Myh6*-Cre^+/–^, *Kcna1*^+/–^). *Myh6*-Cre^+/–^, *Kcna1*^+/–^ mice were generated by crossing hemizygous transgenic *Myh6*-Cre^+/–^ mice with heterozygous *Kcna1*^+/–^ mice. Control mice included the following genotypes from the breeding crosses: *Myh6*-Cre^−/−^; *Kcna1*^+/+^ (wildtype; WT); *Myh6*-Cre^−/−^; *Kcna1*^fl/-^ (fl/-); and *Myh6*-Cre^+/–^; *Kcna1*^+/+^ (Cre). Myh6-Cre mice (Agah *et al*., 1997) were purchased from Jackson Labs (Bar Harbor, MN) where they are cataloged as B6N.FVB(B6)-Tg(Myh6-cre)2182Mds/J (JAX 018972). *Kcna1*^+/–^ mice, which are maintained on a Tac:N:NIHS-BC genetic background, carry a null (KO) allele of the *Kcna1* gene due to targeted deletion of the entire open reading frame (Smart *et al*., 1998). Generation of the *Kcna1*^fl/+^ mice, which are maintained on a C57BL/6N background, was performed as previously described (Trosclair *et al*., 2020). Mice were housed at 22°C, fed *ad libitum*, and submitted to a 12-h light/dark cycle. For lifespan analysis, both experimental and control mice were monitored for mortality until at least the age of 100 days. All experiments were performed in accordance with National Institutes of Health (NIH) guidelines with approval from the Institutional Animal Care and Use Committees of Louisiana State University Health Sciences Center-Shreveport and Southern Methodist University.

### Genotyping

To identify experimental and control mice, genomic DNA was isolated by enzymatic digestion of tail clips using Direct-PCR Lysis Reagent (Viagen Biotech, Los Angeles, CA, USA). Genotypes were determined by performing PCR amplification of genomic DNA using allele-specific primers. The primer sequences used for amplifying the global KO and floxed *Kcna1* alleles have been described previously (Dhaibar *et al*., 2019; Trosclair *et al*., 2020). For detection of the *Myh6*-Cre transgene, the following primer sequences were used to yield a product of 300 bp for the Cre allele: a *Myh6*-Cre specific forward primer (5’-ATGACAGACAGATCCCTCCTATCTCC-3’) and a *Myh6*-Cre specific reverse primer (5’-CTCATCACTCGTTGCATCATCGAC-3’). As an internal control, the *Myh6*-Cre reaction was multiplexed with a primer pair that yielded a 450-bp amplicon of the WT allele of the *Scn2a* gene: 5’-TGCGAGGAGCTAAACAGTGATTAAAG-3’ and 5’-GGCTCCATTCCCTTATCAGACCTACCC-3’.

### Cardiomyocyte isolation

Atrial and ventricular myocytes were enzymatically isolated from hearts of male and female mice (ages 6-8 weeks). Briefly, mice were intraperitoneally injected with 5000 U/kg heparin (Sigma-Aldrich, St. Louis, MO) and euthanized by cervical dislocation. The heart was quickly removed and mounted on a Langendorff apparatus followed by a 3-min retrograde perfusion with oxygenated (100% O_2_) Ca^2+^-free Tyrode’s solution containing (in mmol/l): 140 NaCl, 5.4 KCl, 0.5 MgC_l2_, 10 glucose, and 10 HEPES (pH 7.4; 37°C). Hearts were then perfused with the same Tyrode’s solution but containing Liberase TH enzymes (0.025 mg/ml; Sigma-Aldrich) and bovine serum albumin (BSA; 1 mg/ml; Sigma-Aldrich). Left atrial and left ventricular tissue was then removed, placed in separate petri dishes and minced. Atrial and ventricular myocytes were dispersed in KB solution containing (in mmol/l): 80 KOH, 40 KCl, 25 KH_2_PO_4_, 3 MgSO_4_, 50 glutamic acid, 20 taurine, 1 EGTA, 10 glucose, and 10 HEPES (pH 7.2 with KOH; 20–22°C). Cells were stored at room temperature (20-22°C) for at least 1 h before use. All chemicals used to make the solutions for cell isolations were obtained from Sigma-Aldrich.

### Whole-cell patch-clamp recordings

Whole cell patch-clamp recordings were performed at 37 °C. Borosilicate glass pipette (Warner Instruments, Hamden, CT) microelectrodes were used with tip resistances of 2-3 MΩ when filled with pipette solution. Electrodes were connected to a MultiClamp 700B microelectrode amplifier equipped with a CV-7B head stage (Axon Instruments, Molecular Devices, San Jose, CA). Electrical signals were sampled at 4 kHz and digitized with an Axon analog/digital converter (Digidata 1550B). Data acquisition and analysis were performed using Clampfit software (version 11.4.2, Axon Instruments, Molecular Devices). Dendrotoxin-K (10 nM; Sigma-Aldrich) was used to selectively block Kv1.1 channels, as done previously (Si *et al*., 2018, 2024; Trosclair *et al*., 2021). All chemicals used to make the bath and pipette solutions for recordings were obtained from Sigma-Aldrich. For current-clamp recordings, action potentials were evoked by electrical stimulation with 1-ms, 2-nA current pulses at a frequency of 1 Hz. The bath solution contained (in mmol/l): 126 NaCl, 5.4 KCl, 1.8 CaC_l2_, 1.0 MgCl_2_, 20 HEPES, and 11 glucose (pH=7.4 with NaOH). The pipette solution contained (in mmol/l): 90 K-aspartate, 30 KCl, 10 NaCl, 5.5 glucose, 1.0 MgCl_2_, 10 EGTA, 4.0 Na-GTP, and 10 HEPES (pH=7.2 with KOH).

### Non-invasive electrocardiography

Non-invasive ECG recordings were acquired using the ECGenie system (Mouse Specifics Inc., Framingham, MA). Mice were individually placed on the recording platform and allowed to acclimate for 10 minutes prior to data collection. All recordings were conducted in non-anesthetized, unrestrained animals under controlled environmental conditions (e.g., consistent light levels, temperature, and minimal noise). Following acclimation, ECG signals were recorded for 15-20 minutes via paw contact with the platform electrodes. For each animal, ten recording segments were randomly selected for analysis. Each segment consisted of 20 consecutive, artifact-free ECG waveforms. Ensemble waveforms were then generated from each 20-beat segment using EzCG software (Mouse Specifics Inc.), which also automatically calculated key ECG parameters: heart rate (HR), PR interval, QRS duration, and QT interval (uncorrected and corrected for HR). The corrected QT interval (QTc) was automatically calculated by the software according to a modified Bazett’s formula QTc = QT/ sqrt(RR/100)^1/2^ (Mitchell *et al*., 1998). To obtain mean values for each animal, values from the ten ensemble waveforms (representing a total of 200 beats) were averaged. All automated measurements were visually inspected for accuracy before inclusion in the final dataset.

### Simultaneous electroencephalography-electrocardiography recordings

Mice (4-6 weeks old) of both sexes were anesthetized with an anesthetic cocktail and then surgically implanted with bilateral silver wire electroencephalography (EEG) and electrocardiography (ECG) electrodes (0.005-inch diameter) attached to a microminiature connector (Omnetics Connector Corporation, Minneapolis, MN) for recording in a tethered configuration, as described previously (Mishra *et al*., 2018). The anesthetic cocktail contained ketamine (80-100 mg/kg), xylazine (6-10 mg/kg), and acepromazine (1-2 mg/kg) and was administered by intraperitoneal (i.p.) injection. When anesthetic cocktail was used, the reversal agent Antisedan (1 mg/kg) was administered by subcutaneous (s.c.) injection immediately following completion of the surgical procedure. Carprofen (4-5 mg/kg, s.c.) was given the day of the surgery and 24 hours post-surgery for analgesia. EEG wires were inserted into the subdural space through cranial burr holes overlying parietotemporal cortex for the recording electrodes and above frontal cortex for the ground and reference electrodes. For ECG, two wires were tunneled subcutaneously on both sides of the thorax and sutured in place to record cardiac activity. Mice were allowed to recover from surgery for 24-48 h before recording simultaneous video and EEG-ECG for ≥ 24 h continuously while the animals were housed in a glass container (40-cm length x 20-cm width x 23-cm height) with access to food and water. Biosignals were band-pass filtered by applying 0.3-Hz high-pass and 200-Hz lowpass filters for EEG and a 3.0-Hz high-pass filter for ECG. Sampling rates were set to 500 Hz for EEG and 2 kHz for ECG. Recordings were captured using Ponemah data acquisition and analysis software (Data Sciences International, St. Paul, MN, USA).

### Analysis of EEG-ECG recordings

Quantification of cardiac features was performed using Ponemah software (Data Sciences International, St. Paul, MN), as described previously (Trosclair *et al*., 2020). Heart rate (HR) and heart rate variability (HRV) were estimated from six separate RR interval series derived by sampling 2-min ECG segments every 4 h to provide three light phase (6:00 AM to 6:00 PM) and three dark phase (6:00 PM to 6:00 AM) measurements. The HR and HRV values for each animal were then averaged from the six total segments. RR intervals were only sampled during times when the mouse was stationary and when the ECG showed no abnormalities such as skipped or ectopic beats occurred. Skipped heart beats (e.g., atrioventricular conduction blocks or sinus pauses) were identified from the ECG waveforms over the entire 24-h recording period and defined as a prolongation of the RR interval equaling ≥ 1.5 times the previous RR interval.

### Intracardiac pacing

*In vivo* pacing studies were performed in 4-month old mice of both sexes, as done previously (Glasscock *et al*., 2015; Watts *et al*., 2021). In brief, mice were anesthetized with isoflurane (2% for induction and 1.5% for maintenance of anesthesia; Apollo Tech 3 Vaporizer; NorVap) and placed in a supine position with limbs taped onto surface ECG electrodes of a temperature controlled procedure platform (Rodent Surgical Monitor, Indus Instruments, USA) which maintained core body temperatures at 37.0 ± 0.5 °C. A Millar 1.1F octapolar EP catheter (EPR-800; Millar Instruments) was inserted via an incision in the internal right jugular vein. The catheter was advanced to the right atrium and ventricle using electrogram guidance and pacing capture to verify intracardiac position. A computer-based data acquisition system (PowerLab 16/30; ADI Instruments) was used to record a 2-lead body surface ECG and up to 4 intracardiac bipolar electrograms (LabChart Pro software, version 7; AD Instruments). To induce atrial arrhythmias, programmed electrical stimulation (PES) was performed by delivering 2 ms current pulses to the right atria using an external stimulator (STG-3008 FA; Multi Channel Systems), as done previously (Watts *et al*., 2021). Atrial fibrillation (AF) was defined as the occurrence of rapid and fragmented atrial electrograms with irregular AV nodal conduction and ventricular rhythm for a least 1 s. To induce ventricular arrhythmias, PES was performed by delivering 2-msec current pulses at 400 μA to the right ventricle. As done previously (Trosclair *et al*., 2021), a burst pacing protocol with eight 50-ms and four 30-ms cycle length train episodes was used. This sequence was repeated twenty times every 3 seconds to result in 70.4 seconds of total stimulation time. Ventricular arrythmia (VA) was defined as a sequence of rapid spontaneous ventricular depolarizations following programmed electrical stimulation that lasted ≥1 sec. Each animal first underwent three PES trials to test for AF, followed by three PES trials to test for VA. An animal was considered to have AF or VA if they were detected during any of the three trials.

### Echocardiography

Cardiac ultrasound was performed on isoflurane-anesthetized male and female mice approximately 4 months of age using a VisualSonics Vevo 770 Micro-Imaging System (VisualSonics, Inc., Toronto, ON, Canada) with a 30-MHz transducer, as done previously (Trosclair *et al*., 2021). Two-dimensional directed M-mode transthoracic echocardiographic images along the parasternal short axis were recorded to measure and compare the following parameters between genotypes: left ventricular internal diameter during systole (LVIDs); left ventricular internal diameter during diastole (LVIDd); left ventricular anterior and poster wall thickness during systole (LVADs and LVPWs, respectively); and left ventricular anterior and poster wall thickness during diastole (LVADd and LVPWd, respectively). These parameters were then used to calculate ejection fraction and fractional shortening using standard formulas (Gao *et al*., 2011).

### Flurothyl-induced seizure susceptibility

Flurothyl-induced seizure susceptibility was performed using the same procedure as reported previously (Vanhoof-Villalba et al., 2018). Briefly, mice (postnatal days 30–31) were placed in an air-tight Plexiglas chamber (18.4 × 15.0 × 34.5 cm) and allowed to acclimate for 5 min before being exposed to the liquid convulsant flurothyl (2,2,2-trifluoroethyl ether; Sigma-Aldrich, St Louis, MO, USA). Flurothyl was infused (20 μL min−1) with a syringe pump onto Whatman filter paper (Cytiva, Marlborough, MA, USA) suspended at the top of the chamber from which it was vaporized. The latency (in seconds) was measured as the time from the first drip of flurothyl to the first myoclonic jerk (focal seizure) and to the onset of running and bouncing and/or falling (generalized tonic-clonic seizure). Immediately following generalized seizure onset, the mice were quickly removed to fresh air and placed in a cage for observation and recovery. Each mouse was tested individually and received only one exposure to flurothyl. The testing chamber was cleaned and aerated in between animals.

### Statistical analysis and blinding

Data are presented as mean ± standard deviation (SD). Prism 10 for Windows (GraphPad Software Inc, La Jolla, CA, USA) was used for statistical analyses. Survival curves were evaluated using the Kaplan-Meier log-rank (Mantel-Cox) test. For comparisons involving only two groups, unpaired two-tailed Student’s t tests were employed. For comparisons involving more than two groups, one-way analysis of variance (ANOVA) was used followed by Tukey post-hoc tests. To compare the incidence of pacing-induced arrhythmias and flurothyl-induced mortality across the genotypic groups, individual contingency tables were constructed to reflect the number of animals with and without the appropriate condition in each group. A Chi-square test was performed to assess differences in proportions between groups. For sample sizes less than five, a Fisher’s exact test was used instead to ensure accurate *P* values. Results were considered significant if *P* < 0.05. Biosignal recordings and analyses were performed in a blinded fashion by naming animals or files in a manner that obscured genotype.

### Data availability

The data that support the findings of this study are available from the corresponding author upon reasonable request.

## RESULTS

### 1. Generation and molecular characterization of cardiac-specific *Kcna1* cKO mice

We generated cardiac-specific *Kcna1* cKO mice (i.e., *Myh6*-Cre^+/-^; *Kcna1^del/^*^-^) using the breeding scheme described in the Materials and Methods. The *Myh6*-Cre driver selectively targets cardiomyocytes in the atria and ventricles (Agah *et al*., 1997; Gaussin *et al*., 2002). To confirm tissue-specific deletion of *Kcna1*, we performed PCR using genomic DNA from the brain and heart tissues of cKO and control (WT and fl/-) mice. In cKO mice, PCR analysis of regionally dissected heart tissues revealed the presence of the *Kcna1^del^* allele in both atria and ventricles, but not in the brain, confirming cardiac-specific gene deletion (**Fig. 1**). PCR also detected the *Kcna1^fl^*allele in cKO mouse hearts due to the presence of non-cardiomyocyte cells where *Myh6*-Cre is not expressed.

**Figure 1.**
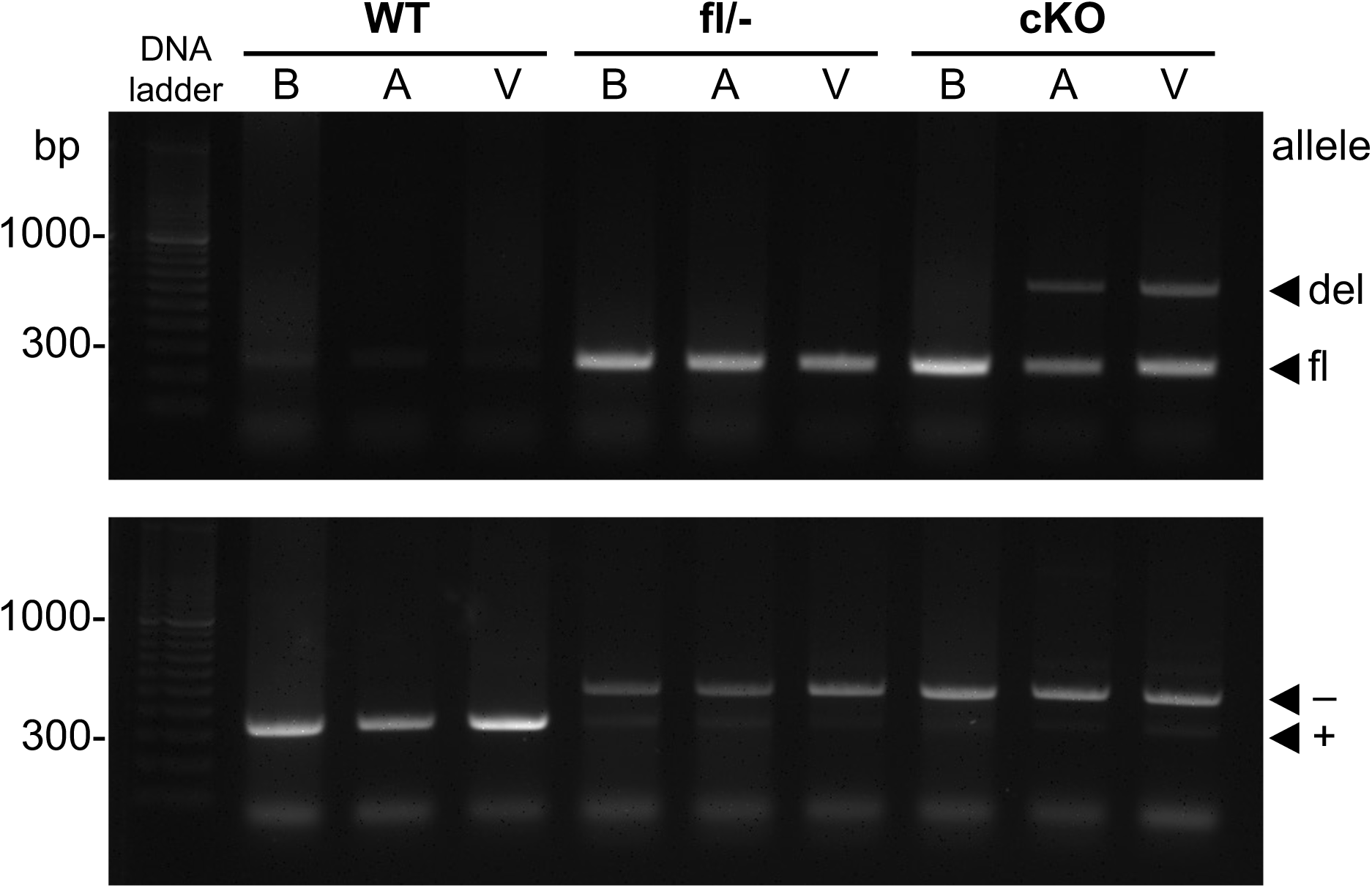
Molecular characterization of cardiac-specific *Kcna1* cKO mice. Representative PCR detection of the *Kcna1* deletion (del; 679 bp), floxed (fl; 260 bp), null (−; 475 bp), and wildtype (WT, +; 337 bp) alleles from genomic DNA isolated from the cortex of the brain (B), atrium (A), and ventricle (V) of WT, *Kcna1*^fl/−^ (fl/-), and cardiac-specific *Kcna1* cKO (cKO) mice.

We next investigated whether cardiac-specific *Kcna1* deficiency causes premature mortality, a phenomenon we previously observed in mice lacking *Kcna1* selectively in either neurons of the brain and spinal cord or in forebrain excitatory neurons (Trosclair *et al*., 2020; Paulhus & Glasscock, 2025). We found that cardiac-specific cKO mice (n=33) exhibited normal lifespans with no premature deaths occurring before four months of age. However, we noted sporadic deaths in both cKO mice and *Myh6*-Cre^+/-^ control animals beginning around six months of age. These deaths were likely attributable to the cardiotoxic effects of the *Myh6*-Cre driver, which have been reported to manifest around this time (Pugach *et al*., 2015). To avoid confounding effects from *Myh6*-Cre-related cardiotoxicity, we limited our experiments to animals four months of age or younger. In addition, our breeding crosses yielded approximately two to three times more female cKO mice than males. As a result, most of our analyses could not be fully sex-matched due to limited availability of male mice; however, this limitation reflects the inherent sex ratio of our breeding outcomes rather than a selection bias.

### 2. Cardiac-specific Kv1.1 deficiency impairs atrial repolarization

Our previous studies have shown that global *Kcna1* KO mice have significantly prolonged action potentials due to the lack of repolarizing I_Kv1.1_ currents (Si *et al*., 2018, 2024; Trosclair *et al*., 2021). To identify the electrophysiological consequences of cardiac-specific *Kcna1* deficiency at the cellular level, we performed patch-clamp recordings on freshly isolated atrial and ventricular cardiomyocytes from cKO and control mice. In whole-cell current-clamp recordings, atrial cKO cells exhibited significantly prolonged action potential durations at 90% repolarization (APD_90_) compared to WT (Tukey-adjusted *P*=0.0017, 1-way ANOVA) and *Myh6*-Cre^+/-^ (Tukey-adjusted *P*=0.0006, 1-way ANOVA) control cells (**Fig. 2A,B**). However, resting membrane potential was not significantly different between genotypes (*P*=0.59, 1-way ANOVA; **Fig. 2C**).

**Figure 2.**
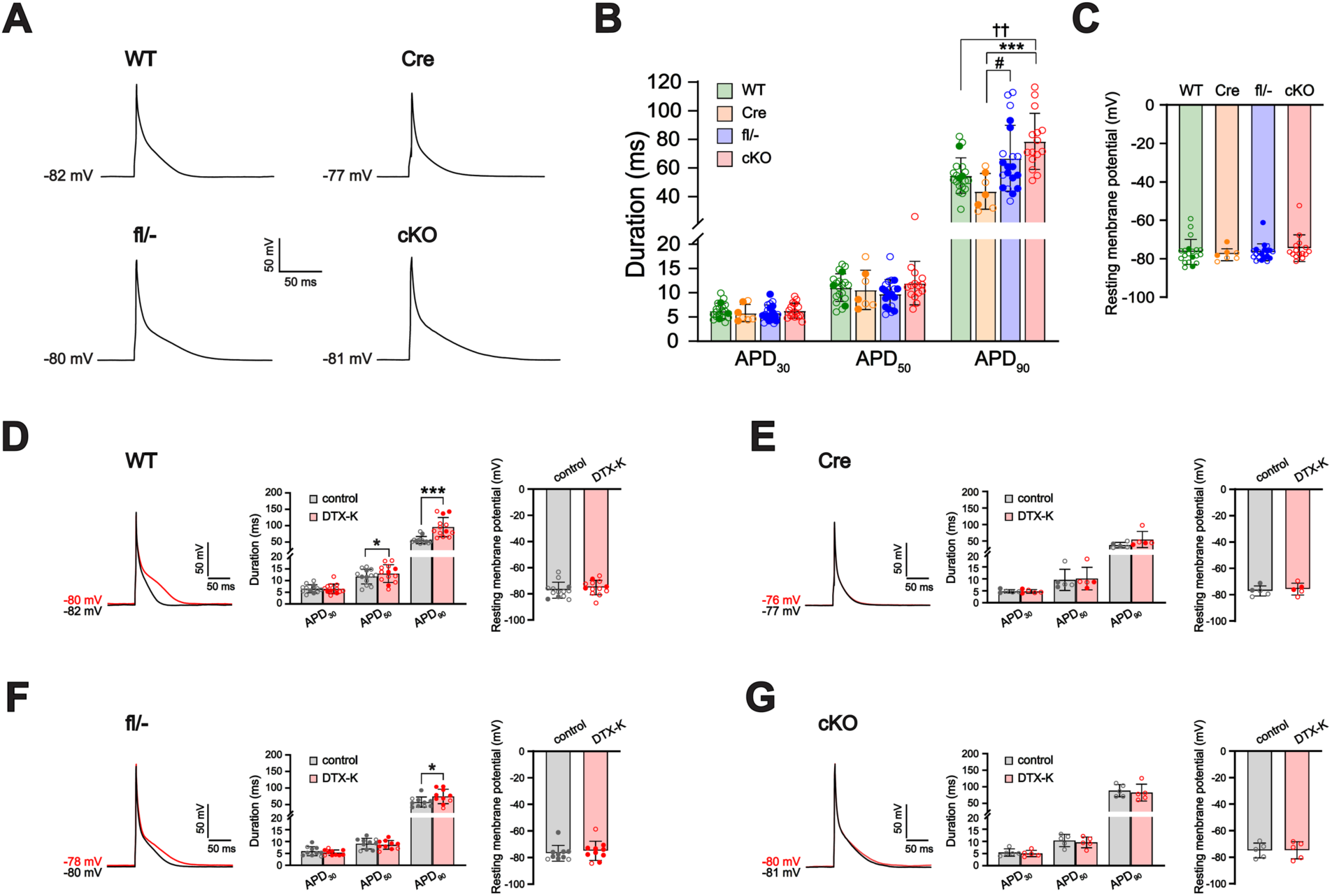
Atrial cardiomyocytes from cardiac-specific *Kcna1* conditional knockout (cKO) mice exhibit prolonged action potential duration (APD). **A,** Representative action potentials recorded in atrial cardiomyocytes from WT, Cre, fl/-, and cKO mice. **B,** Average APD at 30, 50, and 90% repolarization (APD_30_, APD_50_, APD_90_) in WT (n=20/8), Cre (n=7/5), fl/-(n=20/7), and cKO cells (n=15/5). ^††^ adjusted *P* = 0.0017; *** adjusted *P* = 0.0006; ^#^ adjusted *P* = 0.027 (1-way ANOVA; post-hoc Tukey test). **C,** Average resting membrane potential by genotype. *P* = 0.97 (1-way ANOVA). **D-F,** Left panel: Representative action potential recordings in a WT, Cre, fl/-, and cKO cell at baseline (black line) overlaid with the resulting action potential after application of 10 nM DTX-K (red line). Middle and right panels: Average APD_30_, APD_50_, APD_90_ and average resting membrane potential before (control) and after (DTX-K) application of 10 nM DTX-K in WT (n=13/5), Cre (n=5/2), fl/-(n=10/4), and cKO cells (n=5/3). *** *P* < 0.001; * *P* < 0.05 (2-tailed paired Student’s t-test). Sample numbers (n) indicate myocytes per mouse. For the data points, closed circles indicate male sex, and open circles indicate female sex.

Pharmacological inhibition of Kv1.1-containing channels with the Kv1.1-specific blocker dendrotoxin-K (DTX-K) reproduced this effect in both WT and fl/-atrial cells, causing significant prolongation of APD_90_ by 71% (*P*=0.0003, paired Student’s t test) and 29% (*P*=0.019, paired Student’s t test), respectively (**Fig. 2D,F**). In contrast, atrial cardiomyocytes from cKO mice showed no change in action potential duration following DTX-K administration (*P*=0.45, paired Student’s t test), confirming the drug’s specificity for Kv1.1 subunits (**Fig. 2G**).

Consistent with these findings, voltage-clamp recordings showed that DTX-K significantly reduced peak outward K^+^ current density by approximately 24% in WT atrial cells over test potentials from +10 to +50 mV (*P*<0.05, paired Student’s t-test), indicating a contribution of Kv1.1-containing channels to repolarizing currents (**Fig. 3A,C**). This DTX-K-mediated reduction in outward K^+^ currents was absent in atrial cells from cKO mice at the same membrane potentials (**Fig. 3B,D**).

**Figure 3.**
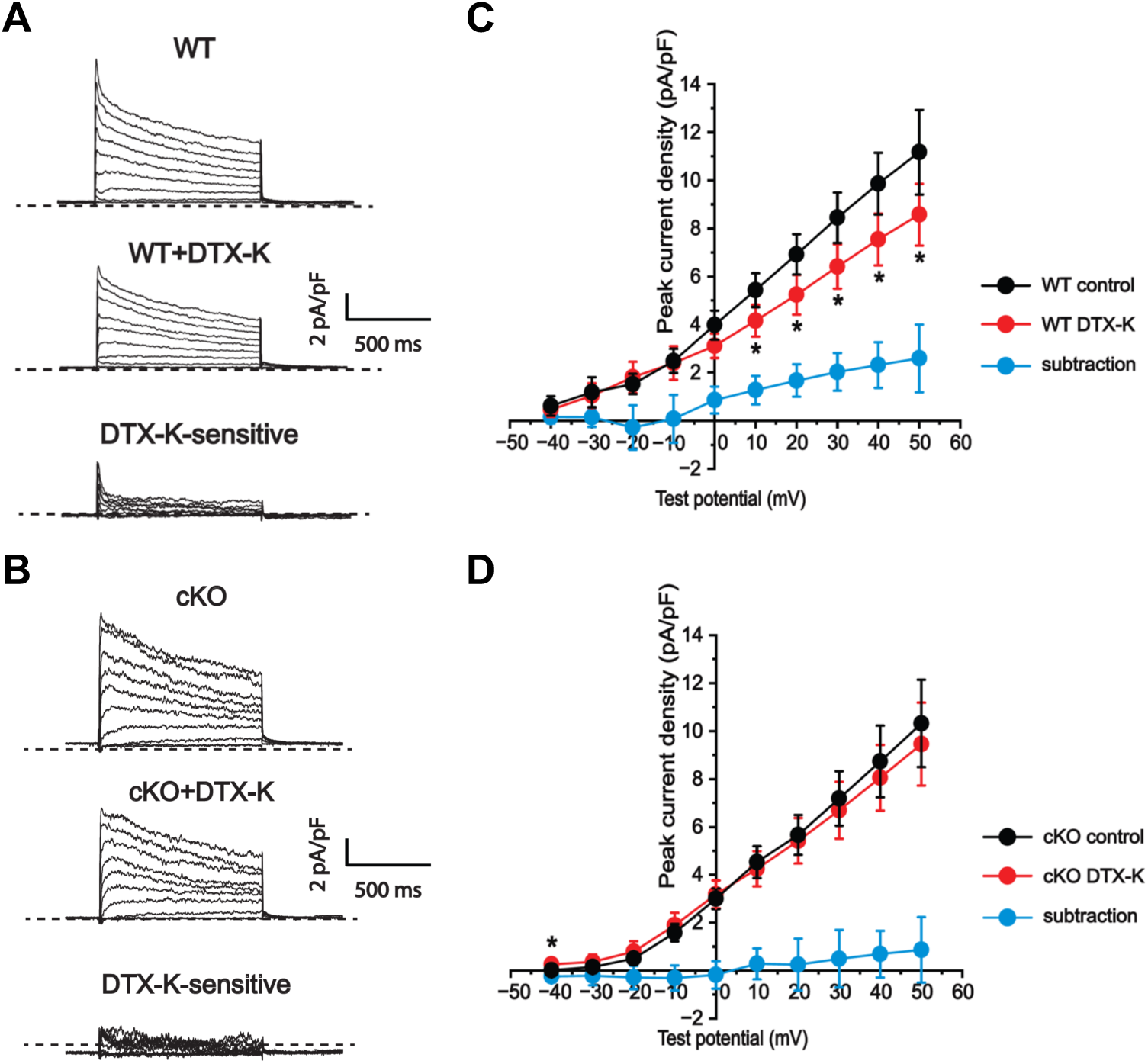
Dendrotoxin-K (DTX-K) reduces outward K^+^ currents in wildtype (WT) atrial cardiomyocytes. **A-B,** Representative raw current traces in WT and cKO cells in response to 1-s depolarizing voltage steps of +10 mV from a holding potential of -50 mV to +50 mV: at baseline (top panel); with application of 10 nM DTX-K (middle panel); and following subtraction of the current difference to isolate the DTX-K-sensitive component (bottom panel). The dotted line in the traces indicates 0 pA. **C-D,** Quantification of the peak current densities before (control) and after (DTX-K) application of DTX-K in WT (n=4/2) and cKO atrial cardiomyocytes (n=7/5), with the blue line (subtraction) indicating the DTX-K-sensitive component. Sample numbers (n) indicate myocytes per mouse. * *P* < 0.05 (2-tailed paired Student’s t-test).

In contrast to atrial cells, ventricular cardiomyocytes from cKO mice exhibited no significant change in action potential duration or resting membrane potential compared to WT cells (**Fig. 4A-C**). However, application of DTX-K caused a small (8-21%) but significant prolongation of APD_90_ in ventricular cardiomyocytes from control mice of all genotypes (**Fig. 4D-F**), suggesting the functional presence of Kv1.1 in the ventricles under normal conditions. In cKO ventricular cardiomyocytes, DTX-K produced no significant effect on APD (**Fig. 4G**).

**Figure 4.**
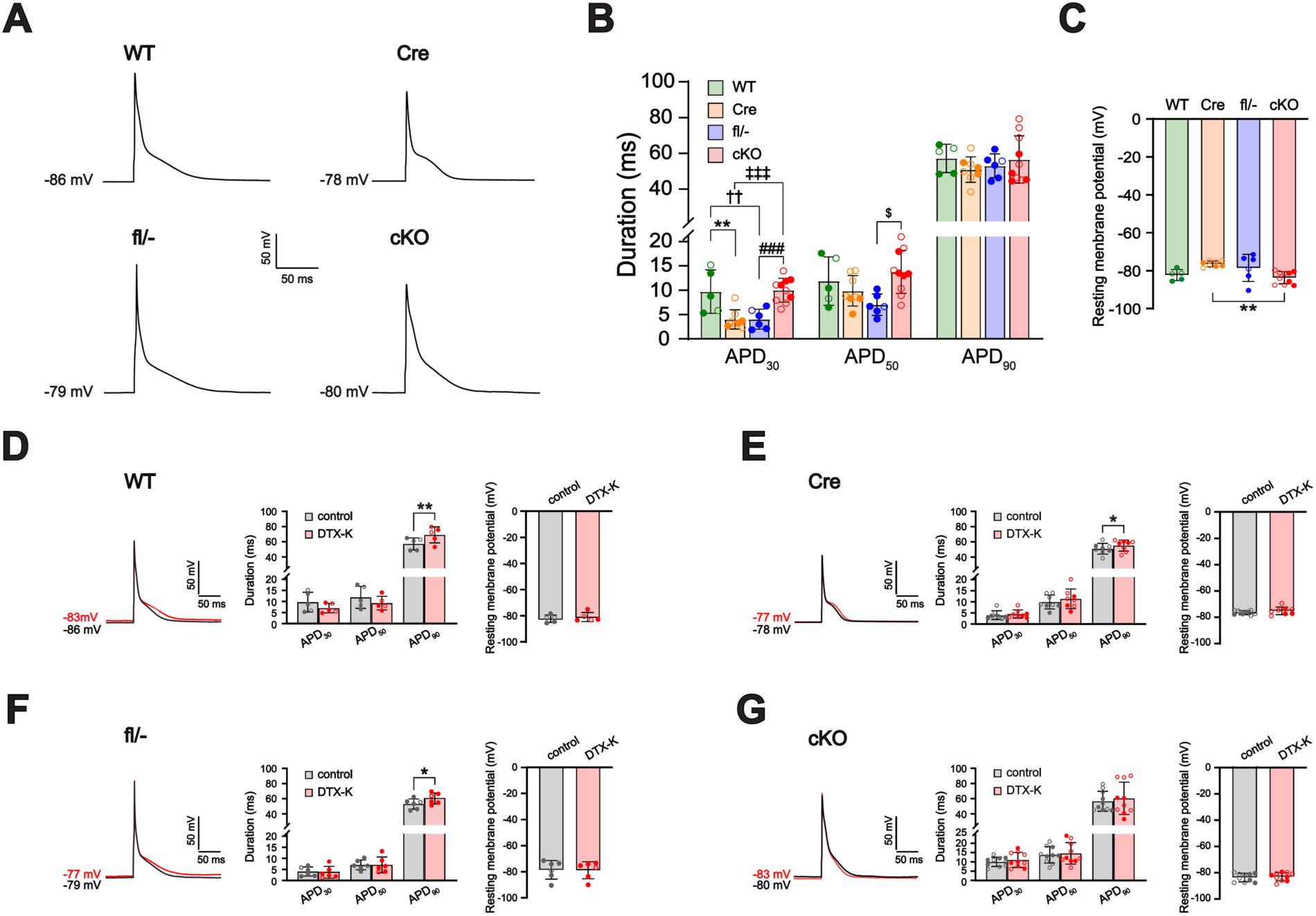
Ventricular cardiomyocytes from cardiac-specific *Kcna1* conditional knockout (cKO) mice exhibit normal action potential duration (APD). **A,** Representative action potentials recorded in ventricular cardiomyocytes from WT, Cre, fl/-, and cKO mice. **B,** Average APD at 30, 50, and 90% repolarization (APD_30_, APD_50_, APD_90_) in WT (n=5/2), Cre (n=8/5), fl/-(n=6/5), and cKO cells (n=10/4). ^‡‡‡^ adjusted *P* = 0.0005; ^###^ adjusted *P* = 0.0014; ^††^ adjusted *P* = 0.0099; ** adjusted *P* = 0.0054 (1-way ANOVA; post-hoc Tukey test). **C,** Average resting membrane potential by genotype. ** adjusted *P* = 0.0041 (1-way ANOVA; post-hoc Tukey test). **D-F,** Left panel: Representative action potential recordings in a WT, Cre, fl/-, and cKO cell at baseline (black line) overlaid with the resulting action potential after application of 10 nM DTX-K (red line). Middle and right panels: Average APD_30_, APD_50_, APD_90_ and average resting membrane potential before (control) and after (DTX-K) application of 10 nM DTX-K in WT (n=5/2), Cre (n=8/5), fl/-(n=6/5), and cKO cells (n=10/4). ** *P* < 0.01; * *P* < 0.05 (2-tailed paired Student’s t-test). Sample numbers (n) indicate myocytes per mouse. For the data points, closed circles indicate male sex, and open circles indicate female sex.

Previous studies have shown that the *Myh6*-Cre driver selectively targets atrial and ventricular cardiomyocytes, but its expression in sinoatrial node (SAN) cells is not well described. To functionally test whether the SAN is targeted by *Myh6*-Cre, we conducted pilot experiments on SAN myocytes isolated from cKO mice to record their spontaneous action potential firing rates. Whereas global *Kcna1* knockout mice exhibit reduced SAN firing rates, our recordings showed that SAN myocytes from cKO mice (n=6 cells) maintained normal firing rates of 331 ± 84 bpm, consistent with healthy control animals in our previous studies (347 ± 90 bpm). This finding confirms that the *Myh6*-Cre driver does not substantially affect Kv1.1 function in SAN cells in cKO mice.

### 3. Cardiac-specific *Kcna1* cKO mice display normal ECG characteristics

To evaluate whether the selective absence of Kv1.1 channels in cardiomyocytes affects *in vivo* cardiac waveform properties, we conducted ECG recordings using both non-invasive and invasive methods. Non-invasive recordings are advantageous for quickly assessing heart rate (HR) and ECG waveform parameters, while invasive recordings, which involve surgically implanted ECG electrodes ideal for longer-term measurements, are more effective for measuring heart rate variability (HRV), an indicator of autonomic influence on the heart.

In non-invasive ECG recordings, cKO mice showed normal HR, which was comparable to that of control mice. Additionally, ECG waveform characteristics exhibited no significant differences between cKO and control animals (**Fig. 5A**, **Table 1**). For invasive recordings, we implanted mice with subcutaneous ECG electrodes, along with EEG brain electrodes to confirm the absence of seizure activity. The animals were then monitored continuously for 24 hours. Consistent with the non-invasive data, the invasive ECG recordings revealed no significant difference in HR between cKO and control mice (**Fig. 5B**, **Table 2**). Furthermore, EEG recordings in cKO mice did not show any obvious pathological brain activity in the form of spontaneous seizures (**Fig. 5B**). We also estimated HRV over the 24-hour recording period (**Table 2**) and found no significant differences in the standard deviation of RR intervals (SDNN), the coefficient of variance (CV), or the root mean square of successive RR intervals (RMSSD). These metrics provide time-domain measures of total autonomic activity (SDNN, CV) and parasympathetic tone (RMSSD). Finally, we quantified the frequency of skipped heart beats, which could be due to either atrioventricular conduction blocks or sinus pauses. Although global KO mice exhibit significant increases in skipped heart beats compared to WT (Glasscock *et al*., 2010), in cKO mice, no differences were apparent (**Table 2**). Overall, these findings indicate that cardiac-specific cKO mice exhibit normal *in vivo* heart function at the level of the ECG.

**Figure 5.**
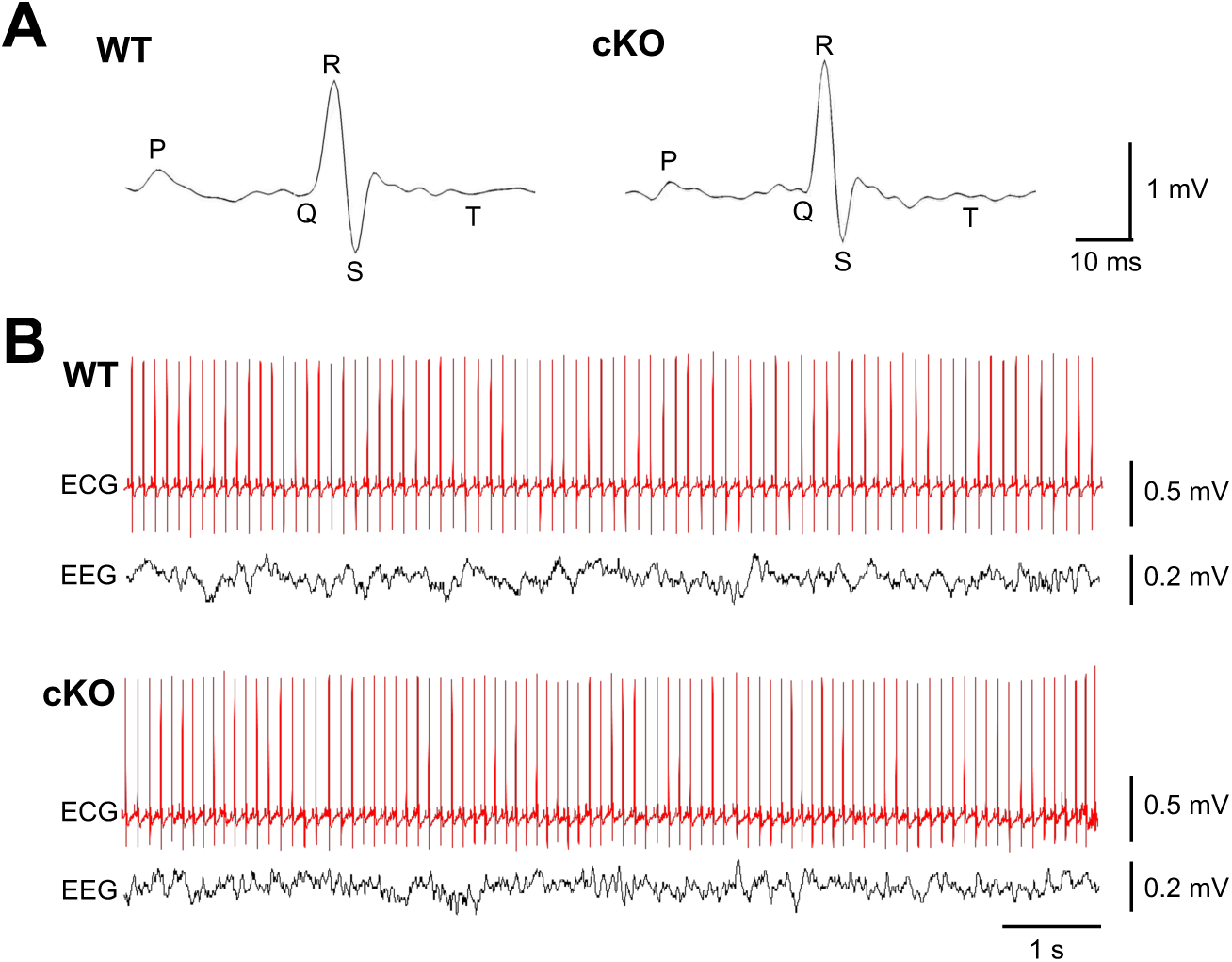
Electrocardiography (ECG) recordings in cardiac-specific *Kcna1* cKO mice. **A,** Representative averaged ensemble ECG waveforms from WT and cKO mice, with each trace showing the mean of 20 individual waveforms recorded non-invasively. The major ECG components (P, Q, R, S, T waves) are labeled. **B,** Representative invasive ECG waveforms recorded simultaneously with electroencephalography (EEG), which appeared normal. For both animals, the EEG trace shows electrical activity in the left temporoparietal lobe.

**Table 1.** ECG waveform characteristics.

|  | **WT**<br>(n=6; 3 M, 3 F) | **Cre**<br>(n=12; 4 M, 8 F) | **fl/-**<br>(n=3; 2 M, 1 F) | **cKO**<br>(n=6; 0 M, 6 F) | P-value |
| --- | --- | --- | --- | --- | --- |
| Heart rate (beats/min) | 766 ± 24 | 730 ± 43 | 722 ± 42 | 704 ± 86 | 0.26 |
| RR (ms) | 79.0 ± 2.7 | 83.7 ± 5.8 | 84.6 ± 5.8 | 87.6 ± 13.3 | 0.31 |
| PQ (ms) | 20.7 ± 0.7 | 21.3 ± 1.5 | 20.3 ± 1.2 | 21.1 ± 1.4 | 0.63 |
| PR (ms) | 26.1 ± 0.7 | 26.5 ± 1.8 | 25.7 ± 1.8 | 26.6 ± 2.4 | 0.87 |
| QRS (ms) | 8.5 ± 0.4 | 8.6 ± 0.8 | 8.6 ± 0.7 | 8.6 ± 1.4 | 0.99 |
| QT (ms) | 38.1 ± 1.2 | 40.0 ± 2.6 | 40.7 ± 2.4 | 41.7 ± 6.3 | 0.37 |
| QTc (ms) | 42.9 ± 0.9 | 43.7 ± 1.4 | 44.3 ± 1.3 | 44.4 ± 3.2 | 0.50 |

Data are expressed as mean ± standard deviation. P-values were derived from 1-way analysis of variance used to compare intragroup differences. The samples sizes indicate the total number of mice measured and the number of males (M) and females (F) in each dataset. QTc is the rate corrected QT value calculated using a modified Bazett's formula.

**Table 2.** Heart rate variability measures.

|  | **WT**<br>(n=5; 2 M, 3 F) | **Cre**<br>(n=6; 4 M, 2 F) | **fl/-**<br>(n=3; 2 M, 1 F) | **cKO**<br>(n=9; 0 M, 9 F) | P-value |
| --- | --- | --- | --- | --- | --- |
| Heart rate (beats/min) | 569 ± 45 | 538 ± 33 | 542 ± 27 | 554 ± 39 | 0.57 |
| SDNN (ms) | 10.5 ± 2.4 | 10.9 ± 2.3 | 8.4 ± 4.3 | 10.4 ± 2.9 | 0.66 |
| CV (%) | 10.0 ± 2.2 | 9.6 ± 1.7 | 7.5 ± 3.4 | 9.6 ± 2.8 | 0.56 |
| RMSSD (ms) | 4.9 ± 0.9 | 6.6 ± 1.9 | 4.1 ± 3.0 | 5.1 ± 1.6 | 0.19 |
| Skipped beats (/hr) | 0.4 ± 0.1 | 2.2 ± 1.2 | 0.5 ± 0.2 | 0.3 ± 0.1 | 0.11 |

Data are expressed as mean ± standard deviation. P-values were derived from 1-way analysis of variance used to compare intragroup differences. The samples sizes indicate the total number of mice measured and the number of males (M) and females (F) in each dataset. Abbreviations: SDNN, standard deviation of beat-to-beat differences; CV, coefficient of variance; RMSSD, root mean square of successive beat-to-beat differences.

### 3. Cardiac-specific *Kcna1* cKO mice exhibit relatively normal arrhythmia susceptibility

In intracardiac pacing studies, global *Kcna1* KO mice exhibit altered susceptibility to atrial fibrillation (AF) and ventricular arrhythmias (VA). To determine if cardiac-specific Kv1.1 deficiency affects arrhythmia susceptibility, we used intracardiac burst pacing stimulation to induce AF and VA, which we monitored using surface ECG and atrial and ventricular electrograms. In response to atrial stimulation, 27% of cKO mice (n=11) developed AF, compared to 0-10% in control mice (n=9-11), but this difference was not statistically significant (**Fig. 6A**). In response to subsequent ventricular stimulation, 9% of cKO mice experienced VA, which was not significantly different from the 20-36% incidence observed in control mice (**Fig. 6B**). These results suggest that the absence of Kv1.1 specifically in the heart does not significantly impact baseline susceptibility to cardiac arrhythmias.

**Figure 6.**
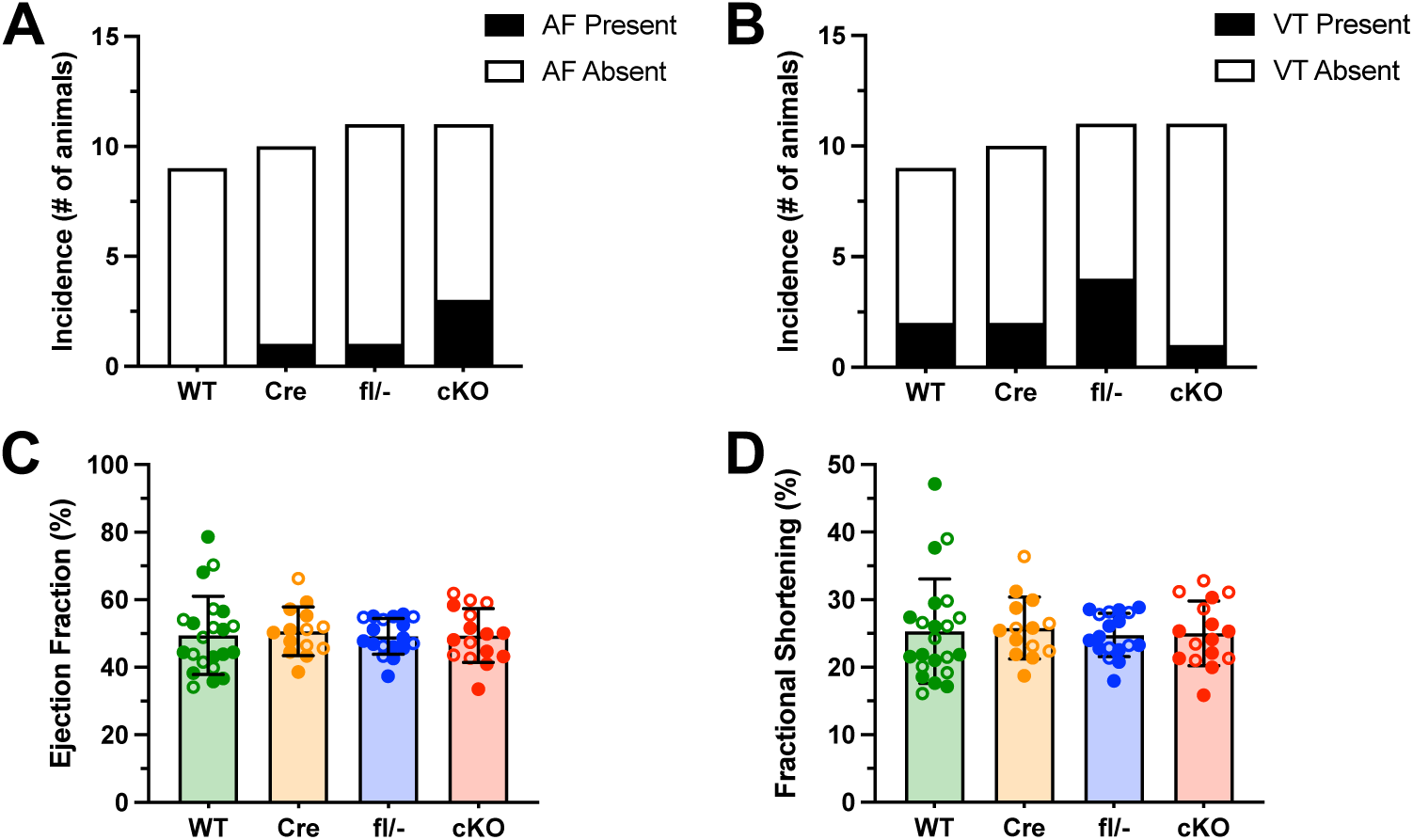
Cardiac-specific *Kcna1* conditional knockout (cKO) mice have normal arrhythmia susceptibility and ventricular contractility and efficiency. In response to programmed electrical stimulation, WT (n=9; 8 M, 1 F), Cre (n=10; 7 M, 3 F), fl/-(n=11; 6 M, 5 F), and cKO (n=11; 7 M, 4 F) mice showed no significant differences in the incidence of (**A**) atrial fibrillation (AF; Chi-square, *P* = 0.41) or (**B**) ventricular arrhythmias (VA; Chi-square, *P* = 0.49). The filled black bars indicate the number of animals that exhibited arrhythmia, and the open white bars show the number of animals that did not exhibit arrhythmia. In echocardiographic imaging, WT (n=22; 12 M, 10 F), Cre (n=14; 9 M, 5 F), fl/-(n=18, 12 M, 6 F), and cKO (n=16; 9 M, 7 F) mice showed no significant differences in (**C**) ejection fraction (1-way ANOVA, *P* = 0.97) or (**D**) fractional shortening (1-way ANOVA, *P* = 0.96). For the data points, closed circles indicate male sex, and open circles indicate female sex.

### 4. Cardiac-specific *Kcna1* cKO mice exhibit normal ventricular contractility and efficiency

Previous studies using *in vivo* echocardiography have shown that global *Kcna1* KO mice display deficits in ventricular function, such as reduced cardiac output and contractility, without obvious structural changes to the heart. Therefore, we used echocardiography to assess ventricular function in cardiac-specific cKO mice and controls. Echocardiographic measurements revealed no significant differences in ejection fraction or fractional shortening between cardiac-specific *Kcna1* cKO mice and control mice (**Fig. 6C,D**), suggesting cardiac-specific Kv1.1 deficiency does not affect overall cardiac output or contractility, respectively.

### 5. Cardiac-specific Kv1.1 deficiency does not alter seizure susceptibility or mortality

Global *Kcna1* KO mice exhibit spontaneous seizures and reduced seizure thresholds when exposed to chemical convulsants. While cardiac-specific cKO mice do not exhibit spontaneous seizures, the lack of Kv1.1 in the heart could potentially increase risk of seizure-related death if a seizure were to occur. To investigate this possibility, we induced seizures in cKO and control mice by exposing them to the volatile convulsant flurothyl, a standard method that reliably elicits an initial focal seizure marked by a myoclonic jerk, followed by a generalized tonic-clonic seizure characterized by running and bouncing. The latency to each seizure type serves as an established measure of seizure threshold. Upon flurothyl exposure, cardiac-specific cKO mice exhibited seizure latencies that were similar to control animals, suggesting their seizure susceptibility was not significantly altered (**Fig. 7A,B**). Furthermore, flurothyl-induced seizures resulted in tonic hindlimb extension followed by death in 29% of cKO mice, which was not significantly different from the 0-17% observed in control mice (**Fig. 7C**). These findings suggest that the lack of Kv1.1 specifically in the heart does not substantially increase the susceptibility to seizure-related death.

**Figure 7.**
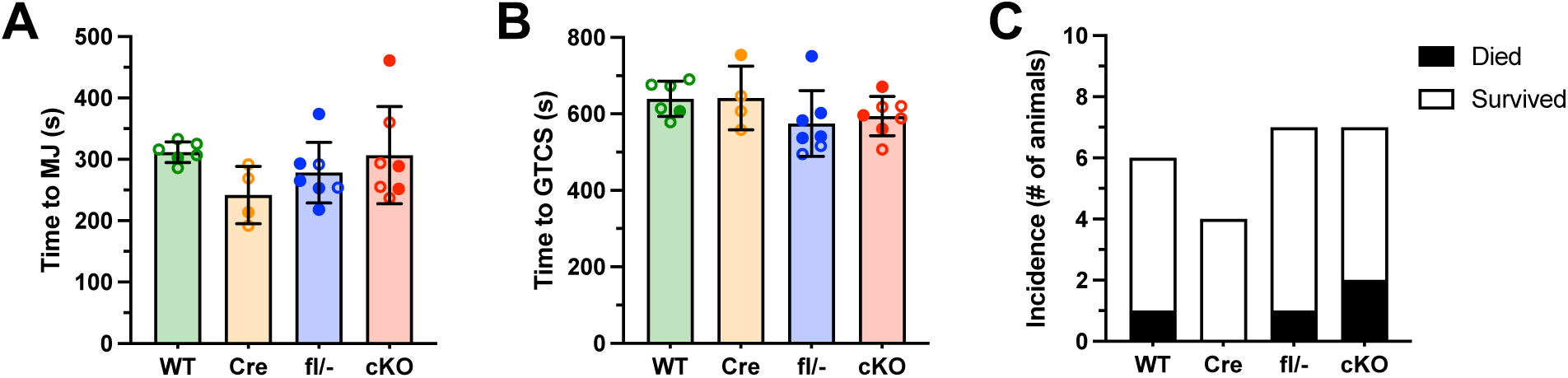
Cardiac *Kcna1* conditional knockout (cKO) mice show normal seizure susceptibility and without increases in seizure-induced mortality. When exposed to flurothyl to induce seizures, WT (n=6; 1 M, 5 F), Cre (n=4; 1 M, 3 F), fl/-(n=7; 5 M, 2 F), and cKO (n=7; 3 M, 4 F) mice exhibited similar latencies to (**A**) myoclonic jerk (MJ) focal seizures (1-way ANOVA, *P* = 0.21) and to (**B**) generalized tonic-clonic seizures (GTCS; 1-way ANOVA, *P* = 0.27), suggesting no significant differences in seizure threshold. **C,** After GTCS onset and subsequent tonic hindlimb extension, cKO mice exhibited no significant differences in the incidence of seizure-associated mortality (Fisher’s Exact, *P* = 0.89). For the data points, closed circles indicate male sex, and open circles indicate female sex.

## DISCUSSION

Our findings show that Kv1.1 plays a heart-intrinsic role in shaping cardiac action potentials, particularly by regulating repolarization in atrial cells. In cardiac-specific *Kcna1* cKO mice, atrial APD was markedly prolonged, whereas ventricular APD remained relatively normal. Because these repolarization deficits were observed in mice lacking Kv1.1 only in the heart, they are not a consequence of epilepsy or seizure-related remodeling of cardiac ion channel expression. Despite impaired cellular repolarization, cKO mice exhibited normal ECG waveform characteristics *in vivo*. Heart rate variability, arrhythmia susceptibility, and measures of cardiac contractility and efficiency, which are altered in global Kv1.1 knockout mice (Glasscock *et al*., 2015; Mishra *et al*., 2017; Trosclair *et al*., 2021), remained normal in cKOs, suggesting that these functional deficits in global KOs arise from Kv1.1 loss in the nervous system rather than from heart-intrinsic Kv1.1 channel deficiency. Finally, cKO mice displayed normal seizure susceptibility and no substantial increase in seizure-induced mortality, implying that loss of Kv1.1 in the heart alone does not appreciably increase the risk of seizure-related death.

Previous work in global Kv1.1 knockout mice and human tissue has demonstrated that Kv1.1 contributes broadly to cardiac electrophysiology and performance. In mice, Kv1.1 protein is expressed in the atria, ventricles, and SAN, with highest abundance in the atria (Glasscock *et al*., 2015; Trosclair *et al*., 2021; Si *et al*., 2024). In humans, Kv1.1 is also detected in both atrial and ventricular tissue, though at more comparable levels between chambers (Glasscock *et al*., 2015; Trosclair *et al*., 2021). Global Kv1.1 loss in mice prolongs action potentials in atrial, ventricular, and SAN myocytes, slows SAN firing rates, alters atrial and ventricular arrhythmia susceptibility, increases parasympathetic tone, and reduces myocardial contractile performance (Glasscock *et al*., 2010, 2015; Mishra *et al*., 2017; Si *et al*., 2018, 2024; Trosclair *et al*., 2021). These cellular electrophysiological changes can be reproduced in WT cells with the Kv1.1-specific blocker DTX-K, which also significantly reduces outward K⁺ current density, confirming a functional role for Kv1.1-containing channels in cardiomyocytes (Si *et al*., 2018, 2024; Trosclair *et al*., 2021). In human atrial tissue, DTX-K-sensitive currents are present and are increased, along with Kv1.1 protein levels, in chronic atrial fibrillation, supporting a role for Kv1.1 in the pathophysiology of persistent AF (Glasscock *et al*., 2015).

Our cardiac-specific *Kcna1* cKO model refines this picture, showing that Kv1.1’s heart-intrinsic role is more limited than suggested by global KO studies. In cKOs, action potential prolongation was confined to atrial myocytes, with normal ventricular repolarization, preserved ECG parameters, stable autonomic function, and intact contractile performance. Interestingly, DTX-K prolonged APD in WT ventricular cardiomyocytes, indicating that Kv1.1-containing channels are normally present in ventricles. The absence of APD changes in cKO ventricles may reflect compensatory channel expression remodeling that masks latent defects; however, prior studies have not detected significant mRNA expression changes in ion channels critical for ventricular repolarization when Kv1.1 is absent (Trosclair *et al*., 2021). Taken together, these findings indicate that the broader systemic cardiac phenotypes of global KO mice require Kv1.1 loss in the nervous system, rather than arising solely from intrinsic deficiency in the heart.

Our study contained some technical limitations that should be considered when interpreting the results. Although we observed no echocardiographic differences between groups, these measurements were performed in younger mice (4 months) compared with 6-8-month old mice in our previous study, which revealed cardiac deficits (Trosclair *et al*., 2021). This age difference was dictated by our experimental workflow, which prioritized reducing animal use by performing echocardiography on the same mice subsequently used for programmed electrical stimulation to assess arrhythmia susceptibility, a terminal procedure. Because 4 months is an optimal age for programmed electrical stimulation and the age used in our prior work, we age-matched to those studies at the expense of performing echocardiography at a younger time point, when heart failure is less likely to manifest. In addition, this younger age was selected to avoid the cardiotoxic effects associated with the *Myh6*-Cre driver, which typically become apparent around 6 months of age (Pugach *et al*., 2015). In our previous work, global KOs only showed differences in ventricular arrhythmia susceptibility following isoproterenol administration (Trosclair *et al*., 2021). In the current study, we did not administer isoproterenol because global KOs show increased atrial fibrillation susceptibility without it (Glasscock *et al*., 2015). Importantly, atrial and ventricular arrhythmia assessments were performed consecutively in the same session, with atrial testing first, so both were conducted without isoproterenol. As a result, we cannot determine whether cKOs exhibit altered ventricular arrhythmia susceptibility under β-adrenergic stimulation. Despite these caveats, the data still support our central conclusion that restricting Kv1.1 deficiency to the heart produces atrial repolarization defects at the cellular level without significantly compromising overall cardiac performance.

In the global Kv1.1 KO model of SUDEP, mortality risk may reflect a pathological interaction between impaired brain and heart function. Consistent with this, we have previously shown that global Kv1.1 KO mice exhibit reduced brain-heart association and connectivity in simultaneous EEG-ECG recordings, suggesting a breakdown in neurocardiac regulation (Mishra *et al*., 2017; Hutson *et al*., 2020). In epilepsy, seizures are often associated with a spectrum of clinical cardiac abnormalities collectively termed epileptic heart syndrome, including atrial fibrillation and/or ventricular arrhythmias, ECG and repolarization changes, altered HRV, myocardial dysfunction detectable by echocardiography, and accelerated atherosclerosis (Verrier *et al*., 2020; Verrier & Schachter, 2024). Our findings indicate that cardiac-specific Kv1.1 deficiency is not sufficient to trigger these broader cardiac abnormalities or to substantially increase seizure-induced mortality. In contrast, other cKO models that restrict Kv1.1 loss to the nervous system are sufficient to cause SUDEP, although death rates are lower than in global KOs, which lack Kv1.1 in both brain and heart (Trosclair *et al*., 2020; Paulhus & Glasscock, 2025). This pattern suggests that Kv1.1 deficiency in the heart, while not fatal or particularly deleterious on its own, may amplify susceptibility to seizure-related death when combined with neural deficiency. One possible implication is that ion channel variants expressed in both brain and heart (Glasscock, 2014; Yu *et al*., 2023), such as Kv1.1, could contribute to SUDEP risk even if they do not produce overt cardiac dysfunction, perhaps through subtle cellular impairments that interact with seizure-induced stress (Bleakley *et al*., 2020). Given the extensive interconnections between the nervous and cardiovascular systems (Slater *et al*., 2024; Goh *et al*., 2025; Fialho & Lin, 2025), these observations support a brain-heart dyssynergy model of SUDEP, in which combined neural and cardiac impairments, even subclinical, disrupt brain-heart coordination and lower the threshold for fatal events.

